# The glare illusion in individuals with schizophrenia

**DOI:** 10.1101/2025.02.23.639792

**Authors:** Hideki Tamura, Aiko Hoshino

**Affiliations:** Department of Computer Science and Engineering Toyohashi University of Technology; Graduate School of Medicine Nagoya University

**Keywords:** schizophrenia, visual illusion, the glare illusion, human psychophysics, brightness

## Abstract

Individuals with schizophrenia are known to demonstrate unique reactions to visual illusions, and prior research has indicated a potential link between their increased susceptibility to geometric illusions and specific symptom profiles. While various illusory experiences have been examined among individuals with schizophrenia, their responses to brightness-related illusions remain poorly understood. In this study, we investigated how individuals with schizophrenia perceive the glare illusion, in which the apparent brightness of the central region is increased. A total of 30 patients with schizophrenia and 34 control participants were recruited. During each trial, a glare or control image (standard stimulus) was presented alongside a control image (comparison stimulus) with one of seven luminance levels. In the glare condition, the standard stimulus was a glare image; in the control condition, two control images were presented, but only the luminance of the comparison stimulus varied. The participants were asked to judge which central region appeared brighter. The results revealed that the individuals with schizophrenia exhibited greater susceptibility to the glare illusion than did the control participants. However, no significant associations were found between susceptibility to the glare illusion and scores assessing symptom severity. These findings suggest that differences in visual processing in patients with schizophrenia may increase their susceptibility to brightness illusions, although this phenomenon is independent of symptom characteristics. Understanding these perceptual alterations may aid in the development of objective measures of visual cognition for patients with schizophrenia.

## 1. Introduction

Visual illusions serve as valuable tools for understanding perception and cognition, as well as the mechanisms of the visual system. Because the processing of visual illusions involves both bottom-up and top-down mechanisms, these illusions enable unique and objective assessments of differences in neural mechanisms. Various visual illusions have been employed to examine differences in visual processing in patients with schizophrenia (Costa, Costa, et al., 2023; King et al., 2017; Kogata & Iidaka, 2018; Notredame et al., 2014). Among these, geometric illusions, motion illusions, and depth inversion illusions have been widely studied. In particular, the Müller–Lyer illusion and facial depth inversion illusion have been shown to be processed differently by individuals with schizophrenia and control participants, suggesting weakened top-down processing in individuals with schizophrenia (Costa, Costa, et al., 2023). Furthermore, previous studies have reported that susceptibility to the Müller–Lyer illusion changes in different stages of schizophrenia (Parnas et al., 2001; Rund et al., 1994). More recently, a study has also demonstrated a relationship between susceptibility to the Müller–Lyer illusion and the stage of schizophrenia (Costa, Barros, et al., 2023), further highlighting the growing interest in illusions as potential biomarkers for early diagnosis and objective assessment.

Thus, numerous studies have investigated the relationships between schizophrenia and various types of visual illusions, including motion illusions, geometric-optical illusions, illusory contours, and depth inversion illusions, on the basis of the classification approach developed by Costa et al. (2023). However, brightness illusions remain largely unexplored in schizophrenia research, despite brightness perception being a fundamental aspect of early visual processing. Previous studies have examined simultaneous brightness contrast in patients with schizophrenia (Grzeczkowski et al., 2018; Kaliuzhna et al., 2019; Yang et al., 2012); however, the findings have been inconsistent. This inconsistency may stem from differences in experimental paradigms or individual variations in contrast sensitivity. Moreover, brightness illusions that rely primarily on local contrast differences and early-stage visual mechanisms may not produce sufficiently strong perceptual effects to differentiate between groups. These findings highlight the need to further investigate brightness illusions that induce stronger perceptual alterations, such as the glare illusion. In contrast to the simultaneous brightness-contrast illusion, the glare illusion increases the perceived brightness beyond what is predicted by local contrast mechanisms, suggesting that additional visual processing mechanisms, possibly involving contextual or top-down modulation, may contribute to this effect. Nevertheless, the perception of brightness illusions in patients with schizophrenia has not yet been thoroughly investigated.

To address this research gap, we investigated how individuals with schizophrenia perceive the glare illusion, a type of brightness illusion. The glare illusion refers to a perceptual phenomenon in which an area appears brighter than its actual physical luminance. This illusion occurs owing to the influence of a surrounding luminance gradient, which increases the perceived brightness of the central region. As illustrated in Figure 1a, when a pattern with a luminance gradient in the surrounding region (glare image) is compared to a pattern without a gradient (control image), the central region of the glare image is perceived as being significantly brighter, despite both images having the same physical luminance level (Agostini & Galmonte, 2002; Suzuki, Minami, & Nakauchi, 2019; Suzuki, Minami, Laeng, et al., 2019; Tamura et al., 2016; Zavagno, 1997). Various geometric patterns have been reported to induce the glare illusion, including squares (Agostini & Galmonte, 2002; Zavagno, 1997; Zavagno et al., 2017), circles (Hanada, 2012; Suzuki, Minami, & Nakauchi, 2019; Tamura et al., 2016), patterns composed of multiple circles (Istiqomah et al., 2022; Kinzuka et al., 2021; Suzuki, Minami, Laeng, et al., 2019), and patterns composed of multiple geometric shapes (e.g., Asahi illusion) (Durand et al., 2024; Laeng et al., 2018; Laeng & Endestad, 2012). Despite their differences, these patterns share a common characteristic: the presence of a surrounding luminance gradient that modulates perceived brightness.

**Figure 1.**
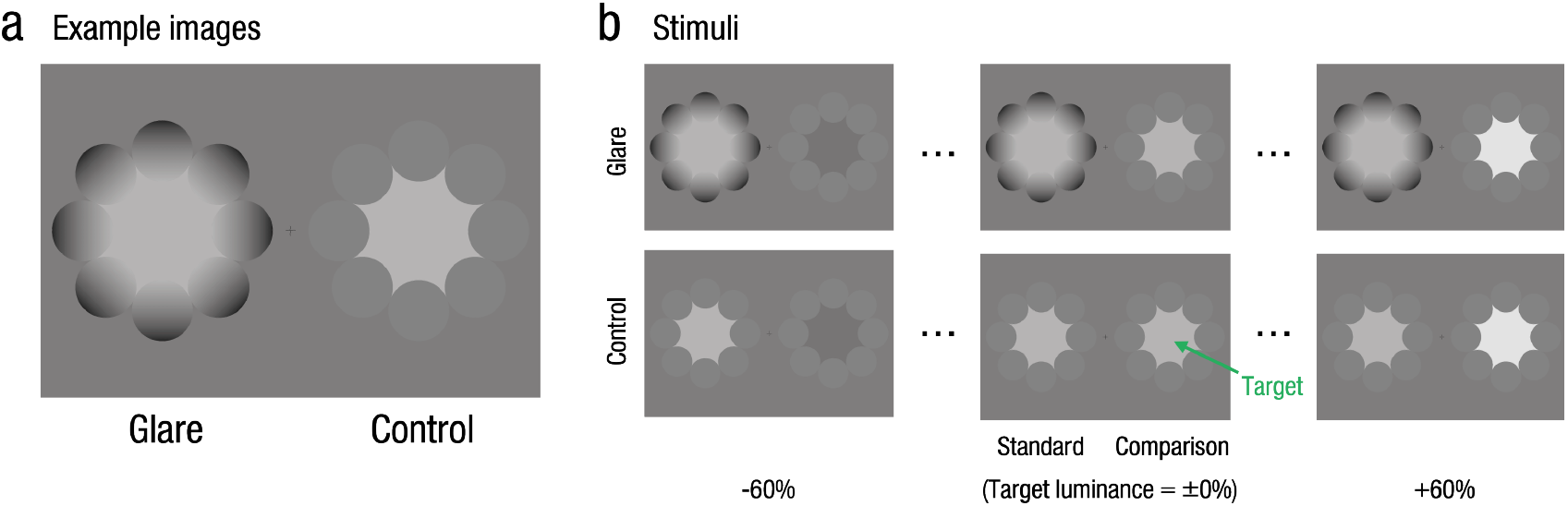
Stimuli. (a) Example images used in the glare illusion experiments. The left image represents a glare image, whereas the right image represents a control image. The luminance values of the central region in both images are identical. (b) Experimental stimuli. The upper row displays the stimuli used in the glare condition, and the lower row shows the stimuli used in the control condition. Examples of luminance pairs at −60%, ±0%, and +60% are shown from left to right.

Moreover, in addition to the increase in the perceived brightness of the central region, the central area has been reported to be perceived as glowing (Tamura et al., 2016) or even dazzling (Hanada, 2012, 2023). In addition to subjective evaluations measured through psychophysical experiments, the physiological responses associated with viewing the glare illusion have been investigated. For example, pupil measurements have shown that pupillary constriction occurs more strongly when the glare illusion is observed (Durand et al., 2024; Istiqomah et al., 2022; Laeng et al., 2018; Laeng & Endestad, 2012; Suzuki, Minami, & Nakauchi, 2019; Suzuki, Minami, Laeng, et al., 2019; Zavagno et al., 2017). Furthermore, a study examining the glare illusion in individuals with autism spectrum disorder (ASD) reported that both individuals with ASD and control participants presented similar subjective increase in brightness and pupillary constriction (Laeng et al., 2018). Given that ASD and schizophrenia have both been associated with atypical visual perception, including differences in sensory integration and perceptual organization (Simmons et al., 2009), it is possible that individuals with schizophrenia may also exhibit altered brightness perception due to differences in visual information processing mechanisms.

Therefore, the aims of this study were to investigate how individuals with schizophrenia perceive the glare illusion and whether their susceptibility to this illusion differs from that of controls. A psychophysical experiment was conducted in which individuals with schizophrenia (SZ) and control participants (CN) were both asked to report which of the two presented images—the glare image or the control image—appeared brighter. The susceptibility to the illusion was then compared between the two groups. Given that individuals with SZ exhibit increased susceptibility to geometric illusions (Costa, Barros, et al., 2023), we hypothesized that their altered visual information processing may lead to increased susceptibility to the glare illusion, a type of brightness illusion. Additionally, we examined whether susceptibility to the glare illusion is correlated with subjective perceptions of brightness in daily life. A relationship between the glare illusion effect and brightness perception in daily life may suggest that individual differences in visual processing, which are influenced by daily perceptual experiences, influence how illusions are perceived. This could provide insights into how contextual factors contribute to the formation of brightness illusions. Furthermore, in the SZ group, we explored whether susceptibility to the glare illusion was associated with the severity of the disease. While previous studies have reported that susceptibility to the Müller–Lyer illusion is influenced by the severity of the disease in individuals with SZ (Costa, Barros, et al., 2023), the relationship between brightness illusions and symptom severity remains unclear. By examining this potential association, we aimed to assess whether individual differences in illusion susceptibility are linked to the clinical characteristics of patients with SZ.

## 2. Methods

### 2.1 Participants

A total of 30 participants with schizophrenia (SZ) and 34 control participants (CN) participated in the experiment. The mean age and standard deviation (SD), as well as the sex ratio, for each group were as follows: SZ group, 52.83 ± 8.6 years, with 70% males; and CN group, 46.38 ± 8.48 years, with 32% males. The sample size was determined to ensure that at least 30 individuals were included in each group. In prior research on SZ and visual illusions (Costa, Barros, et al., 2023), the sample consisted of 14 patients with chronic SZ and 24 controls, indicating that the sample size in this study is reasonable. The SZ group included outpatients with chronic conditions receiving standard treatment, primarily pharmacotherapy, psychotherapy, and rehabilitation. Additionally, the Brief Psychiatric Rating Scale (BPRS) and the Global Assessment of Functioning (GAF) scores of the individuals in the SZ group were measured by the medical staff responsible for each patient. The mean and SD of these scores were 34.77 ± 4.06 and 46.43 ± 12.25, respectively. All procedures were approved by the Research Ethics Committee of the Graduate School of Medicine, Nagoya University, Japan (authorization number: 2023-0406), in 2024, and all ethical standards laid out in the Declaration of Helsinki were followed. All participants provided informed consent prior to participating in the study.

### 2.2 Apparatus

The experimental stimuli were presented on the display of a Surface Pro 8 (2880 × 1920 resolution, 60 Hz refresh rate). The monitor was calibrated using a spectroradiometer (SR-3AR, TOPCON), ensuring a gamma value of 1. The participants observed the tablet display, which was placed on a table, while seated in a chair. They were instructed to maintain a viewing distance of 50 cm from the display. The experiment was conducted in an indoor environment, ensuring that no external light directly illuminated the monitor or the participants. The average illuminance of the display at the beginning of the experiment for all the participant was 598.94 lx. The experimental script was executed using MATLAB R2024a and Psychtoolbox 3.0 (Brainard, 1997; Kleiner et al., 2007; Pelli, 1997).

### 2.3 Stimuli

Figure 1 shows the experimental stimuli, namely, glare illusion patterns composed of eight circles (Kinzuka et al., 2021; Suzuki, Minami, Laeng, et al., 2019). Two types of images were prepared; glare images and control images (Figure 1a). In the glare images, the luminance gradient of the surrounding inducers increased from the outside (darker area, *Y* = 0.74 cd*/*m^2^) to the inside of the image (brighter area, *Y* = 82.09 cd*/*m^2^, defined as the reference luminance). In the control images, the luminance gradient of the inducers was removed and replaced with a uniform luminance equal to the average luminance of the inducer. Thus, the average luminance of the inducers in the glare and control images was identical. The luminance of the central region in the control images was set to seven levels relative to the target luminance of *Y* = 82.09 cd*/*m^2^, corresponding to −60%, −40%, −20%, ±0%, +20%, +40%, and +60%, forming the luminance factor (the −60%, ±0%, +60% examaples are shown in Figure 1b). Each image was presented with an inscribed circle in the central region, subtending the visual angle by 5 degrees.

### 2.4 Procedure

After a fixation point was presented at the center of the screen for 1 second, the stimuli were simultaneously displayed on both sides of the fixation point. In the glare condition, the pair consisted of a glare image (standard stimulus) and a control image (comparison stimulus). In the control condition, the pair consisted of a control image (standard stimulus) and another control image (comparison stimulus). In the standard stimulus, the luminance of the central region was fixed at *Y* = 82.09 cd*/*m^2^; this stimulus was a glare image in the glare condition and a control image in the control condition. The comparison stimulus was always a control image, and the luminance of the central region of this image was set to one of the seven levels defined in by the luminance factor.

The participants, while fixating on the fixation point, compared the central regions of the two images presented and responded by touching the image they perceived as brighter. After the participant provided a response, the next trial began. The stimuli were displayed until a response was provided. The left and right positions of the standard and comparison stimuli were randomized. Each participant completed 112 trials (2 conditions × 7 luminance levels × 2 left–right positions × 4 repetitions), and the trial order was randomized. A break was provided every 28 trials, and the participants were allowed to rest for as long as they needed.

Afterward, the participants completed a questionnaire, which was provided to assess their subjective evaluations of daily life. Among the questions, three items specifically addressed brightness perception, and participants were asked to respond using a 5-point Likert scale: Q1: I find car headlights to be excessively bright; Q2: I find sunlight to be excessively bright; and Q3: My eyes tend to feel fatigued during dim light conditions in the evening. The responses were rated as follows: 5 = strongly agree, 4 = agree, 3 = neutral, 2 = disagree, and 1 = strongly disagree.

### 2.5 Data analysis

Data analysis was conducted using R (version 4.3.3). For each participant, the probability of responding that the comparison stimulus appeared brighter was calculated at each luminance level. The response probabilities across the seven luminance levels for each pattern condition were fitted with a probit function to derive the psychometric functions for each condition. Additionally, the point of subjective equality (PSE) was calculated from these functions. Data that failed to fit the probit function were excluded, as this was considered indicative of participants not performing the task appropriately. As a result, 6 participants from the SZ group and 3 participants from the CN group were excluded. The remaining data from the 24 individuals in the SZ group and 31 participants in the CN group were used for subsequent analyses.

To examine the presence of brightness enhancement effects induced by the glare illusion, the PSE values were predicted using a linear mixed-effects model. The model included the fixed effects of group (G: SZ vs. CN) and condition (C: glare vs. control), the interaction between group and condition with an intercept, and a random effect of the participants. Thus, the model was specified as follows:

*PSE* = *G* + *C* + *G* : *C* + 1 + (1|*P articipantID*). Post hoc pairwise comparisons were conducted with Bonferroni adjustment for p values.

Following the approach of Costa et al. (2023), susceptibility to the illusion was defined as the PSE difference, which was calculated as the difference between the PSE values in the glare and control conditions. This value was computed for all participants, and differences between the SZ and CN groups were tested using Welch’s two-sample t test.

In addition, pairwise correlations among the subjective brightness responses (Q1, Q2, and Q3), participants’ ages, BPRS scores, and GAF scores were analyzed.

## 3. Results

Figure 2 shows the experimental results. Figure 2a shows the PSE values, and the mean PSE values and SD for each group and condition pair were 111.61±39.18 and 82.52±3.68 (glare and control conditions in the SZ group) and 92.87±20.07 and 82.73±1.60 (glare and control conditions in the CN group). The results of the linear mixed-effects models (LMMs; see Table 1) revealed a significant main effect of the stimulus condition, indicating that the glare stimulus was perceived as brighter than the control stimulus was (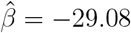, 95% CI [−41.04, −17.13], *t*(53.00) = −4.77, *p <* .001, Cohen’s *d* = −1.38). Additionally, there was a significant main effect of group, with the individuals in the SZ group perceiving the stimuli as brighter than the individuals in the CN group did 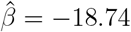, 95% CI [−30.05, −7.43], *t*(105.99) = −3.25, *p* = .002, Cohen’s *d* = −0.89). Furthermore, a significant interaction effect was observed (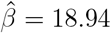, 95% CI [3.02, 34.87], *t*(53.00) = 2.33, *p* = .024, Cohen’s *d* = 0.90).

**Table 1.** Summary of the parameters of the LMMs based on the PSE values.

| term | $\hat{\beta}$ | 95% CI | $t$ | $df$ | $p$ | Cohen’s $d$ |
| --- | --- | --- | --- | --- | --- | --- |
| Intercept | 111.61 | [103.11, 120.10] | 25.75 | 105.99 | < .001 | 5.28 |
| Stimulus | -29.08 | [-41.04, -17.13] | -4.77 | 53.00 | < .001 | -1.38 |
| Group | -18.74 | [-30.05, -7.43] | -3.25 | 105.99 | .002 | -0.89 |
| Stimulus × Group | 18.94 | [3.02, 34.87] | 2.33 | 53.00 | .024 | 0.90 |

**Figure 2.**
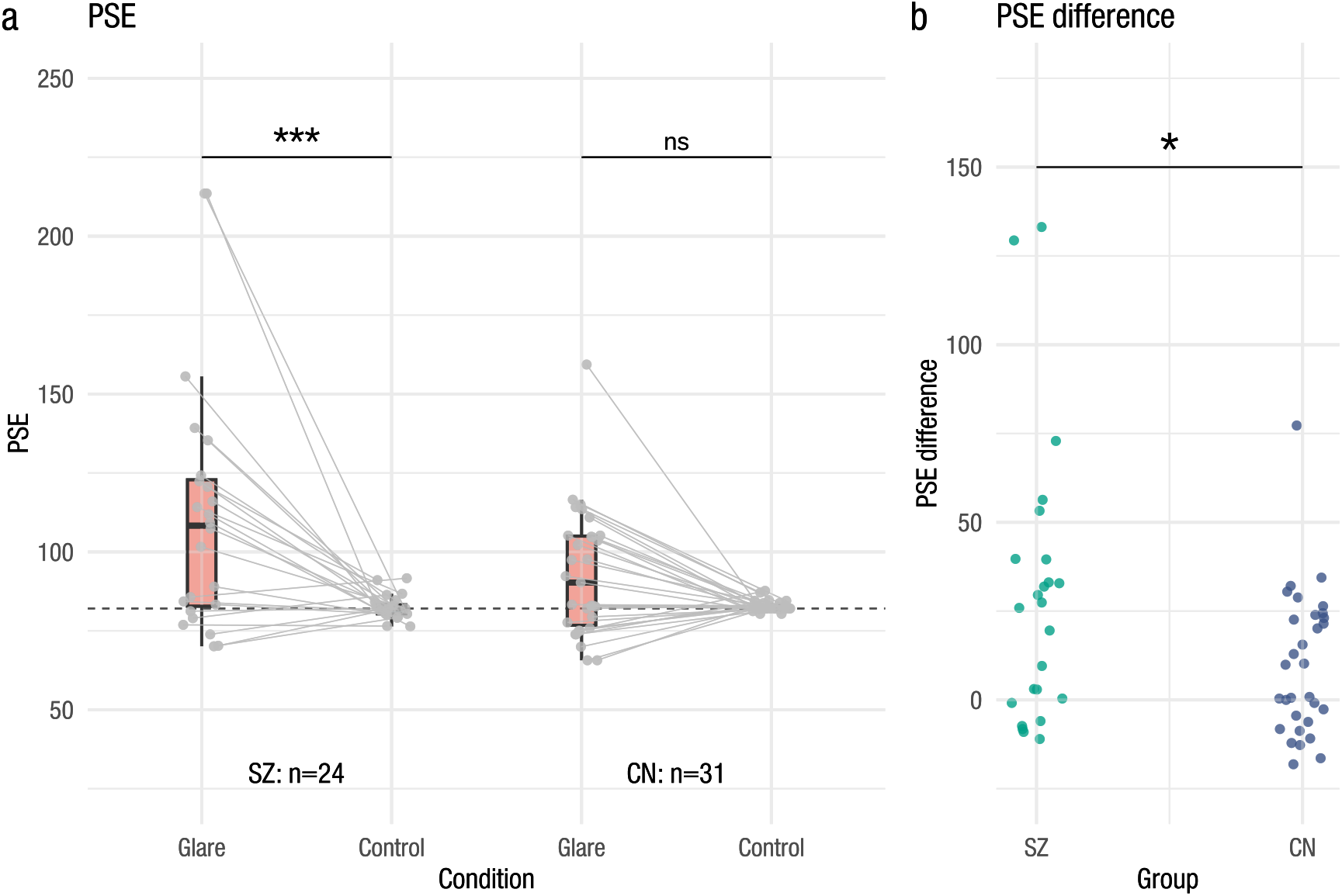
Experimental results. (a) PSE values for each condition. The boxplot displays the distribution, with the data points for individual participants overlaid. The horizontal dashed line represents the target luminance (Y = 82.09 cd/m2). (b) Differences in the PSE values among the groups. Each dot represents the PSE value for an individual participant.

Post hoc tests revealed that the individuals in the SZ group perceived the glare stimuli as brighter than the control stimuli (Δ*M* = 29.08, 95% CI_Bonferroni(6)_ [12.36, 45.80], *t*(53) = 4.77, *p*_Bonferroni(6)_ *<* .001). In contrast, no significant difference between the perception of the glare and control stimuli was observed among the individuals in the CN group (Δ*M* = 10.14, 95% CI_Bonferroni(6)_ [−4.57, 24.85], *t*(53) = 1.89, *p*_Bonferroni(6)_ = .386). The complete comparison results are presented in Table 2.

**Table 2.** Post hoc comparisons with t tests and Cohen’s d values for each pair.

| Contrast | $\Delta M$ | 95% $CI_{\text{Bonferroni}(6)}$ | $t$ | $df$ | $p_{\text{Bonferroni}(6)}$ | Cohen's $d$ |
| --- | --- | --- | --- | --- | --- | --- |
| Glare SZ - Control SZ | 29.08 | [12.36, 45.80]<sup>a</sup> | 4.77 | 53 | < .001<sup>a</sup> | 0.65 |
| Glare SZ - Glare CN | 18.74 | [3.22, 34.26]<sup>b</sup> | 3.25 | 105.99 | .009<sup>b</sup> | 0.31 |
| Glare SZ - Control CN | 28.88 | [13.36, 44.40]<sup>c</sup> | 5.00 | 105.99 | < .001<sup>c</sup> | 0.48 |
| Control SZ - Glare CN | -10.34 | [-25.86, 5.18]<sup>d</sup> | -1.79 | 105.99 | .456<sup>d</sup> | -0.17 |
| Control SZ - Control CN | -0.20 | [-15.73, 15.32]<sup>e</sup> | -0.04 | 105.99 | > .999<sup>e</sup> | 0.00 |
| Glare CN - Control CN | 10.14 | [-4.57, 24.85]<sup>f</sup> | 1.89 | 53 | .386<sup>f</sup> | 0.26 |

Figure 2b shows the susceptibility of the individuals to the glare illusion (PSE difference), which was significantly greater among the individuals with SZ than among the individuals in the CN group (Δ*M* = 18.94, 95% CI [1.12, 36.76], *t*(32.58) = 2.16, *p* = .038, Cohen’s *d* = 0.61). This finding indicates that individuals with SZ were more likely to perceive the glare illusion as brighter than were the participants in the CN group.

As shown in Figure 3, no significant correlations were observed between the PSE difference and the results of the subjective questionnaire for all participants (PSE difference and Q1: *r*(53) = −.048, 95% CI [−.309, .220], *p* = .727; PSE difference and Q2: *r*(53) = .085, 95% CI [−.184, .343], *p* = .536; PSE difference and Q3: *r*(53) = .049, 95% CI [−.219, .310], *p* = .723). Similarly, no significant correlation was found between the PSE difference and age of the participants (*r*(53) = .168, 95% CI [−.102, .415], *p* = .219). Additionally, among the individuals with SZ, there were no significant correlations between the PSE difference and the BPRS or GAF scores (PSE difference and BPRS: *r*(22) = −.042, 95% CI [−.438, .368], *p* = .847; PSE difference and GAF: *r*(22) = .224, 95% CI [−.197, .575], *p* = .293). On the other hand, significant correlations were observed among the responses on the three subjective questionnaire items among all participants (Q1 and Q2: *r*(53) = .591, 95% CI [.386, .740], *p <* .001; Q2 and Q3: *r*(53) = .434, 95% CI [.190, .627], *p <* .001; and Q1 and Q3: *r*(53) = .310, 95% CI [.049, .532], *p <* .05).

**Figure 3.**
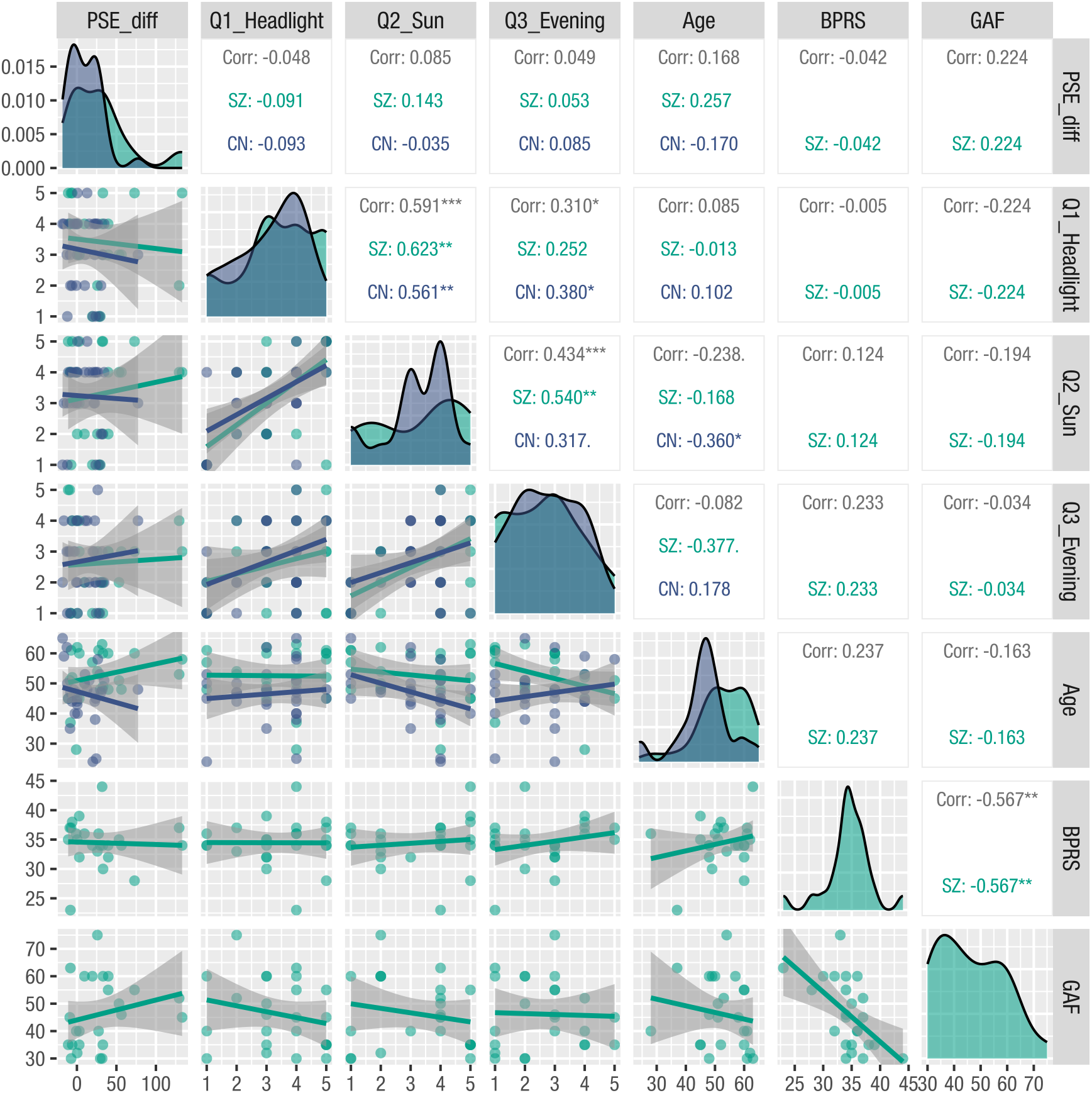
Distribution and correlation plots among pairs of variable. The diagonal matrix displays the distributions of the PSE difference, Q1 response, Q2 response, Q3 response, age, BPRS score, and GAF score for each group. The lower triangular matrix presents scatter plots of variable pairs, including regression lines and their confidence intervals. The upper triangular matrix shows the correlation coefficients between variable pairs (from top to bottom: overall, SZ group only, and CN group only). The asterisks and dots indicate significant differences:. < .1, * < .05, ** < .01, *** < .001.

## 4. Discussion

In this study, we investigated the perception of the glare illusion in individuals with schizophrenia (SZ) using a psychophysical experiment. A comparison of the PSE values confirmed the increase in brightness effect consistently demonstrated in previous glare illusion studies, with glare images being perceived as brighter than control images. However, this effect was observed only among the individuals with SZ, whereas no significant difference between the perceived brightness of the glare and control images was observed in the control (CN) group. A primary contribution of this study is the finding that susceptibility to the glare illusion (PSE difference) was significantly greater among individuals with SZ than among the participants in the CN group. This finding suggests that altered visual information processing in individuals with SZ may influence brightness perception. However, susceptibility to the glare illusion was not found to be correlated with subjective brightness perception in daily life, age, or the severity of the disease. These results indicate that while individuals with SZ were more likely to perceive the glare image as brighter than the control image, this increased susceptibility was not associated with the severity of their symptoms. This finding indicates that the ways in which SZ affects visual function may vary between geometric illusions (e.g., Costa, Barros, et al., 2023) and brightness perception.

One possible explanation for the increased brightness perception among the individuals with SZ is that cumulative alterations in visual processing, such as differences in contrast sensitivity and sensory gating, may lead to increased susceptibility to the glare illusion. In other words, cognitive disorganization and reduced cognitive flexibility, which are characteristic of SZ, may have contributed to the heightened susceptibility of these individuals to visual illusions. Considering previous theories on the mechanisms underlying the glare illusion, differences in either bottom-up or top-down processing may have led to a compensatory shift in the reliance on the other mechanism, making individuals with SZ more prone to perceiving the illusion as brighter. First, a reduction in bottom-up processing efficiency (Luo et al., 2021) could increase reliance on top-down processing, leading to greater susceptibility to the illusion. Specifically, individuals with SZ exhibit differences in the early stages of visual processing, such as reduced contrast sensitivity (Kantrowitz et al., 2009) and altered sensory gating responses (Ludewig et al., 2003; Preuss et al., 2011). These differences may result in less effective visual information processing, thereby increasing the influence of higher-order perceptual processes, such as the top-down interpretation of light diffusion on the basis of past experiences, ultimately leading to increased susceptibility to brightness illusions. SZ is also associated with differences in top-down processing (Berkovitch et al., 2018; Silverstein et al., 2006), which may increase reliance on bottom-up processing when the glare illusion is observed. In this case, the luminance gradient of the inducing stimulus, a key element of the illusion (Hanada, 2012, 2023), might have strongly influenced low-level visual processing, further increasing the effects associated with the illusion. Additionally, studies on brightness induction have yielded conflicting findings regarding the visual mechanisms involved in brightness perception (retina, lateral geniculate nucleus, and primary visual cortex) in patients with SZ. While some studies have reported relatively typical processing (Kinoshita & Komatsu, 2001; Rossi & Paradiso, 1999; Yang et al., 2012), other studies have suggested variations in these mechanisms (Delord et al., 2006). Given these inconsistencies, further research is needed to determine how alterations in bottom-up and top-down processing contribute to brightness perception in patients with SZ. Understanding these mechanisms may provide insights into the broader differences in visual perception associated with this disorder.

Although a main effect of pattern type (glare vs. control) was observed across all participants, an analysis within the CN group did not reveal a significant difference between the perception of the glare and control images (Figure 2a, right panel). This finding differs from previous studies on subjective brightness perception in the glare illusion, which consistently reported that glare images are perceived as brighter than control images are (e.g., Tamura et al., 2016). One possible explanation for this discrepancy is the age difference between the control group in this study and the participants in previous studies. Specifically, age-related factors may have influenced the perception of the glare illusion. Most previous studies on the perception of the glare illusion have focused primarily on participants in their twenties, whereas the average age of the individuals in the control group was higher in the present study. These findings suggest that aging may modulate or diminish the brightness enhancement effect associated with the glare illusion.

This study has several limitations that should be considered. First, while the present study focused solely on behavioral responses, pupillary responses may also play a role in the perception of such illusions, as suggested by previous studies (e.g., Kinzuka et al., 2021; Suzuki, Minami, Laeng, et al., 2019). Laeng et al. reported that pupillary constriction occurred in individuals with ASD while observing the glare illusion (Laeng et al., 2018). Thus, individuals with SZ may exhibit similar pupillary responses to the glare illusion, which should be further explored. Second, while brightness perception was increased among the individuals with SZ, it remains unclear whether the previously reported glow and dazzling perceptions are also affected by SZ symptoms (Tamura et al., 2016). Previous research has examined not only which stimulus appears brighter but also which stimulus appears more dazzling in the glare illusion (Hanada, 2012, 2023). In future studies, new insights could be provided by directly assessing how individuals with SZ perceive glare. Third, given the high heterogeneity of SZ symptoms among patients (Tandon et al., 2013), further studies are needed to explore the relationship between the susceptibility to brightness illusions and other psychiatric conditions.

## 5. Conclusion

In this study, we investigated how individuals with SZ perceive the glare illusion and whether their susceptibility to this illusion differs from that of control participants. The results revealed that individuals with SZ exhibited greater susceptibility to the glare illusion than did the control group. However, unlike susceptibility to geometric illusions, this increased sensitivity was not associated with the severity of the SZ symptoms. Additionally, no significant relationship was found between susceptibility to the glare illusion and subjective brightness perceptions in daily life. These findings suggest that while differences in visual information processing in individuals with SZ may contribute to their increased susceptibility to brightness illusions, this effect does not appear to be directly linked to symptom-related factors or individual differences in brightness perception in daily life. Further research is needed to clarify the mechanisms underlying this brightness enhancement effect in individuals with SZ and to explore how contextual factors influence the perception of brightness illusions.

## Acknowledgments

This work was supported by JSPS KAKENHI (Grant Number JP22K17987) and Tokai Pathways to Global Excellence (T-GEx), which are part of the MEXT Strategic Professional Development Program for Young Researchers. The authors wish to thank Tatsumi Asakura for supporting the data collection.

## Data availability

The data that support the findings of this study are available from the corresponding author upon reasonable request. However, data sharing is subject to restrictions as per the ethical approval obtained for this study, and only deidentified or limited datasets may be shared to ensure participant confidentiality.

## Artificial intelligence

During the preparation of this work, the authors used ChatGPT 4o to improve the language, and the manuscript has been proofread by native English speakers through an English editing service. After using the tool and service, the authors reviewed and edited the content as needed and take full responsibility for the content of the publication.

## References

Agostini, T., & Galmonte, A. (2002). A new effect of luminance gradient on achromatic simultaneous contrast. Psychonomic Bulletin and Review, 9 (2), 264—269. 10.3758/bf03196281

Berkovitch, L., Cul, A. D., Maheu, M., & Dehaene, S. (2018). Impaired conscious access and abnormal attentional amplification in schizophrenia. NeuroImage: Clinical, 18, 835–848. 10.1016/j.nicl.2018.03.010

Brainard, D. H. (1997). The Psychophysics Toolbox. Spatial Vision, 10 (4), 433—6. http://www.ncbi.nlm.nih.gov/pubmed/9176952

Costa, A., Barros, M., Mortari, M., Caixeta, F., & Maior, R. (2023). Stage-dependent sensitivity to Müller-Lyer visual illusion in schizophrenia patients. Behavioural Brain Research, 114173. 10.1016/j.bbr.2022.114173

Costa, A., Costa, D. L., Pessoa, V. F., Caixeta, F. V., & Maior, R.S. (2023). Systematic review of visual illusions in schizophrenia. Schizophrenia Research, 252, 13–22. 10.1016/j.schres.2022.12.030

Delord, S., Ducato, M. G., Pins, D., Devinck, F., Thomas, P., Boucart, M., & Knoblauch, K. (2006). Psychophysical assessment of magno- and parvocellular function in schizophrenia. Visual Neuroscience, 23 (3-4), 645–650. 10.1017/s0952523806233017

Durand, J.-B., Marchand, S., Nasres, I., Laeng, B., & Castro, V.D. (2024). Illusory light drives pupil responses in primates. Journal of Vision, 24 (7), 14. 10.1167/jov.24.7.14

Grzeczkowski, L., Roinishvili, M., Chkonia, E., Brand, A., Mast, F. W., Herzog, M. H., & Shaqiri, A. (2018). Is the perception of illusions abnormal in schizophrenia? Psychiatry Research, 270, 929–939. 10.1016/j.psychres.2018.10.063

Hanada, M. (2012). Luminance Profiles of Luminance Gradients Affect the Feeling of Dazzling. Perception, 41 (7), 791–802. 10.1068/p7070

Hanada, M. (2023). Effects of a gap between the central and surrounding regions with luminance gradients on the feeling of being dazzled. I-Perception, 14 (3), 20416695231176132. 10.1177/20416695231176132

Istiqomah, N., Suzuki, Y., Kinzuka, Y., Minami, T., & Nakauchi, S. (2022). Anisotropy in the peripheral visual field based on pupil response to the glare illusion. Heliyon, 8 (6), e09772. 10.1016/j.heliyon.2022.e09772

Kaliuzhna, M., Stein, T., Rusch, T., Sekutowicz, M., Sterzer, P., & Seymour, K.J. (2019). No evidence for abnormal priors in early vision in schizophrenia. Schizophrenia Research, 210, 245–254. 10.1016/j.schres.2018.12.027

Kantrowitz, J. T., Butler, P. D., Schecter, I., Silipo, G., & Javitt, D. C. (2009). Seeing the World Dimly: The Impact of Early Visual Deficits on Visual Experience in Schizophrenia. Schizophrenia Bulletin, 35 (6), 1085–1094. 10.1093/schbul/sbp100

King, D. J., Hodgekins, J., Chouinard, P. A., Chouinard, V.-A., & Sperandio, I. (2017). A review of abnormalities in the perception of visual illusions in schizophrenia. Psychonomic Bulletin & Review, 24 (3), 734–751. 10.3758/s13423-016-1168-5

Kinoshita, M., & Komatsu, H. (2001). Neural Representation of the Luminance and Brightness of a Uniform Surface in the Macaque Primary Visual Cortex. Journal of Neurophysiology, 86 (5), 2559–2570. 10.1152/jn.2001.86.5.2559

Kinzuka, Y., Sato, F., Minami, T., & Nakauchi, S. (2021). Effect of glare illusion-induced perceptual brightness on temporal perception. Psychophysiology, 58 (9), e13851. 10.1111/psyp.13851

Kleiner, M., Brainard, D. H., Pelli, D. G., Broussard, C., Wolf, T., & Niehorster, D. (2007). What’s new in Psychtoolbox-3? Perception, 36 (14), 1—16. 10.1068/v070821

Kogata, T., & Iidaka, T. (2018). A review of impaired visual processing and the daily visual world in patients with schizophrenia. Nagoya Journal of Medical Science, 80 (3), 317–328. 10.18999/nagjms.80.3.317

Laeng, B., & Endestad, T. (2012). Bright illusions reduce the eye’s pupil. Proceedings of the National Academy of Sciences, 109 (6), 2162–2167. 10.1073/pnas.1118298109

Laeng, B., Færevaag, F. S., Tanggaard, S., & Tetzchner S. von. (2018). Pupillary Responses to Illusions of Brightness in Autism Spectrum Disorder. I-Perception, 9 (3), 2041669518771716. 10.1177/2041669518771716

Ludewig, K., Geyer, M. A., & Vollenweider, F.X. (2003). Deficits in prepulse inhibition and habituation in never-medicated, first-episode schizophrenia. Biological Psychiatry, 54 (2), 121–128. 10.1016/s0006-3223(02)01925-x

Luo, H., Zhao, Y., Fan, F., Fan, H., Wang, Y., Qu, W., Wang, Z., Tan, Y., Zhang, X., & Tan, S. (2021). A bottom-up model of functional outcome in schizophrenia. Scientific Reports, 11 (1), 7577. 10.1038/s41598-021-87172-4

Notredame, C.-E., Pins, D., Deneve, S., & Jardri, R. (2014). What visual illusions teach us about schizophrenia. Frontiers in Integrative Neuroscience, 8, 63. 10.3389/fnint.2014.00063

Parnas, J., Vianin, P., Sæbye, D., Jansson, L., Larsen, A. V., & Bovet, P. (2001). Visual binding abilities in the initial and advanced stages of schizophrenia. Acta Psychiatrica Scandinavica, 103 (3), 171–180. 10.1034/j.1600-0447.2001.00160.x

Pelli, D. G. (1997). The VideoToolbox software for visual psychophysics: transforming numbers into movies. Spatial Vision, 10 (4), 437—442. 10.1163/156856897×00366

Preuss, U. W., Zimmermann, J., Watzke, S., Langosch, J., Siafarikas, N., Wong, J. W. M., Hamm, A., & Weike, A. (2011). Short-Term Prospective Comparison of Prepulse Inhibition between Schizophrenic Patients and Healthy Controls. Pharmacopsychiatry, 44 (03), 102–108. 10.1055/s-0031-1271687

Rossi, A. F., & Paradiso, M. A. (1999). Neural Correlates of Perceived Brightness in the Retina, Lateral Geniculate Nucleus, and Striate Cortex. The Journal of Neuroscience, 19 (14), 6145–6156. 10.1523/jneurosci.19-14-06145.1999

Rund, B. R., Landrø, N. I., Orbeck, A. L., & Nysveen, G. (1994). Mueller-Lyer illusion and size estimation performance in schizophrenics compared to normal controls. Scandinavian Journal of Psychology, 35 (3), 193–197. 10.1111/j.1467-9450.1994.tb00943.x

Silverstein, S., Hatashita-Wong, M., Schenkel, L., Wilkniss, S., Kovács, I., Fehér, A., Smith, T., Goicochea, C., Uhlhaas, P., Carpiniello, K., & Savitz, A. (2006). Reduced top-down influences in contour detection in schizophrenia. Cognitive Neuropsychiatry, 11 (2), 112–132. 10.1080/13546800444000209

Simmons, D. R., Robertson, A. E., McKay, L. S., Toal, E., McAleer, P., & Pollick, F. E. (2009). Vision in autism spectrum disorders. Vision Research, 49 (22), 2705–2739. 10.1016/j.visres.2009.08.005

Suzuki, Y., Minami, T., Laeng, B., & Nakauchi, S. (2019). Colorful glares: Effects of colors on brightness illusions measured with pupillometry. Acta Psychologica, 198, 102882. 10.1016/j.actpsy.2019.102882

Suzuki, Y., Minami, T., & Nakauchi, S. (2019). Pupil Constriction in the Glare Illusion Modulates the Steady-State Visual Evoked Potentials. Neuroscience, 416, 221–228. 10.1016/j.neuroscience.2019.08.003

Tamura, H., Nakauchi, S., & Koida, K. (2016). Robust brightness enhancement across a luminance range of the glare illusion. Journal of Vision, 16 (1), 10. 10.1167/16.1.10

Tandon, R., Gaebel, W., Barch, D. M., Bustillo, J., Gur, R. E., Heckers, S., Malaspina, D., Owen, M. J., Schultz, S., Tsuang, M., Os, J. V., & Carpenter, W. (2013). Definition and description of schizophrenia in the DSM-5. Schizophrenia Research, 150 (1), 3–10. 10.1016/j.schres.2013.05.028

Yang, E., Tadin, D., Glasser, D. M., Hong, S. W., Blake, R., & Park, S. (2012). Visual Context Processing in Schizophrenia. Clinical Psychological Science, 1 (1), 5–15. 10.1177/2167702612464618

Zavagno, D. (1997). Some New Luminance-Gradient Effects. Perception, 28 (7), 835–838. 10.1068/p2633

Zavagno, D., Tommasi, L., & Laeng, B. (2017). The Eye Pupil’s Response to Static and Dynamic Illusions of Luminosity and Darkness. I-Perception, 8 (4), 2041669517717754. 10.1177/2041669517717754

